# Chloroquine attenuates hypoxia-mediated autophagy to curb thrombosis- an *ex vivo* and *in vivo* study

**DOI:** 10.1101/2024.04.23.590850

**Authors:** Propanna Bandyopadhyay, Yash T. Katakia, Sudeshna Mukherjee, Syamantak Majumder, Shibasish Chowdhury, Rajdeep Chowdhury

**Author notes:** Corresponding author: Rajdeep Chowdhury, PhD, Department of Biological Sciences, Birla Institute of Technology and Science (BITS Pilani), Pilani campus, Rajasthan – 333031, address.

## Abstract

**Background:** Hypoxia can trigger the activation of blood platelets, leading to thrombosis. If not addressed clinically, it can cause severe complications and fatal consequences as well. The current treatment regime for thrombosis is often palliative and includes long-term administration of anticoagulants, which has the risk of over-bleeding in injury and other secondary effects as well. This demands a deeper understanding of the process and exploration of an alternative therapeutic avenue. Interestingly, recent studies demonstrate that platelets though atypical and enucleated, possess components of autophagy machinery. This cellular homeostatic process though well-studied in non-platelet cells, is under-explored in platelets.

**Methods:** In this study, we report an activation of autophagy in rat-derived platelets cultured under physiologically relevant hypoxic condition (10% O_2_) *ex vivo*. Furthermore, autophagy was triggered *in vivo* when rats were exposed to hypobaric hypoxic conditions. Subsequently, restriction or surgical ligation of the inferior vena cava (IVC) was performed to induce thrombus formation. Post confirming the impact of autophagy induction on platelet functioning, it was inhibited, and then platelet activation and aggregation status were evaluated using light transmission aggregometry, flow cytometry, immunofluorescence and immunoblotting.

**Results:** Herein, we show that autophagy inhibition with the potent autophagy inhibitor-CQ, a repurposed FDA-approved drug, can significantly reduce platelet activation, both in *ex vivo* and *in vivo* settings. CQ withdrawal reversed the phenomenon indicating a dynamic effect. Thereafter, in flow restriction or surgical ligation model, interestingly, CQ-pre-treated rats showed reduced clotting ability. Importantly, CQ at the stipulated dose was found to be non-toxic to the tissues, as analyzed through histological staining.

**Conclusion:** Thus, we propose that the repurpose of the FDA approved drug CQ can attenuate hypoxia-induced thrombosis through inhibition of autophagy and can be explored as an effective therapeutic alternative.

**Highlights:**

- Exposure of *ex-vivo* cultured platelets to physiologically relevant hypoxic condition (O_2_ concentration) can induce autophagy causing them to get activated and eventually aggregate.
- FDA approved drug chloroquine (CQ) is able to inhibit autophagy in anucleate cellular fragments, platelets, similar to nucleated cells and curb platelet functioning under hypoxic condition in both *ex-vivo* and *in-vivo* model.
- Flow restriction model mimicking deep vein thrombosis, emphasizes on the effect of CQ on impeding thrombus formation when used at a non-toxic dosage.

## Introduction

Platelets are anucleate cellular fragments that essentially function in maintaining hemostasis and vascular integrity. They are the smallest member among all the blood cellular components and actively participate in blood clot formation wherever and whenever necessary.^1^ With a very short life span of 7-10 days in the human body and 4-5 days in rodents, platelet physiology is quite interesting with limited number of cellular organelles, storage vesicles like granules and lysosomes.^2^ Normal platelet functioning involves adhesion, activation, and aggregation of platelets to the site of injury. Accordingly, they also possess several proteins on their outer membrane which facilitate their adhesion to either endothelial cells, neighboring platelets or other circulating blood cells upon inflammation or injury.^3,4^ These platelet aggregates are bound together by fibrin threads to form outer mesh like structure visible over open wounds.^4^ Interestingly, just as platelets start adhering, they change their state from ‘inactive’ to ‘active’ simultaneously changing their morphology from discoid to spherical shape, and extending cytoskeletal filaments to have a stronger grip with the surrounding cells and tissues.^5,6^ However, in their inactive state as well platelets have a number of ongoing cellular processes which help them to maintain homeostasis and keep them upbeat and armed for a necessary transition to the ‘active’ state.

One such homeostatic process of relevance in platelets is autophagy.^7^ Importantly, the discovery of components of autophagy machinery is very recent and subsequent studies further demonstrate its role not only in platelet activation but also in megakaryopoiesis and further also in inflammation and host immune responses.^8^ However, given the versatility of platelet functions and the already identified multifaceted role of autophagy recognized primarily in nucleated cells, there might be multiple cellular functions regulated by autophagy in the atypical platelet cells. Our understanding toward the same is still at its infancy.

Amongst the diverse cellular function of autophagy, the most widely accepted is its active involvement in elimination of damaged or unwanted proteins and organelles *via* engulfment into a vacuolar structure called autophagosome which fuses with lysosomes to degrade the contents. It is also induced majorly under stress conditions like nutrient starvation, ROS generation, and chemotherapeutic stress playing a major role in restoring cellular/tissue balance. ^9,10^ Finally, autophagy and its role in inflammatory states, host immunity, and elimination of intra-cellular pathogens have also dramatically increased in recent years thus integrating its role in cellular defense mechanism. In addition to above, one of the several process where autophagy and its homeostatic machinery is highly implicated is hypoxia. Autophagy aids in cellular redox and nutrient hemostasis under hypoxic conditions. Herein, a cellular hypoxic state can be defined as the disruption in the balance between amount of oxygen required by the cells compared to what is acquired. Physiologically this may lead to oxygen tension build up in the arteries due to inability of lungs to properly oxygenate. This state is often associated with various cardiovascular disorders.^11^ Deep vein thrombosis (DVT) is one such condition that occurs as a low lander travels to a higher altitude and encounters a hypoxic state.^12^ This results in unnecessary clot formation called thrombus, which may initially occur in the calf muscles, but can subsequently travel throughout the body *via* blood vessels resulting in a state known as venous thromboembolism (VTE). Occasionally, such clots can reach vital organs like lungs and heart blocking the movement of blood (pulmonary thromboembolism) causing serious conditions like myocardial infarction and stroke.^13,14^

Importantly, since hypoxia can activate platelets and autophagy as well, in this study we planned to dissect the link between them. Though there are a few studies demonstrating autophagic status in platelets, yet the role of autophagy inhibition on platelet functionality under hypoxic condition has not yet been explored. Our study not only provides insights into the role of autophagy in platelet activation and thrombosis under low oxygen conditions *ex vivo* and *in vivo*, but also offers a novel therapeutic alternative for prevention of hypoxia-induced thrombotic clots.

## Materials and Methods

### Chemicals and reagents

Trisodium citrate (Na_3_C_6_H_5_O_7_ #20242) and Potassium chloride (KCl, #39594) were acquired from s d fine-chem limited. Citric acid (C₆H₈O₇ #22585) was obtained from Fisher Scientific. Sodium chloride (NaCl, #1.93606.5021), Sodium dihydrogen phosphate (NaH_2_PO_4_ #61784505001730) and Diethyl Ether (#1.00923.2521) was procured from Merck. Magnesium chloride (MgCl_2_, #1349130) and thiazolyl blue tetrazolium bromide (MTT, #33611) were purchased from SRL. HEPES sodium salt (#TC066-25G) and Gelatin (#TC041-500G) were purchased from Himedia. Chloroquine (CQ, #C6528), Phenylmethylsulfonyl fluoride (PMSF, #P7626), Thrombin (#T4393-100UN) and Dextrose (C₆H₁₂O₆ #G7021) were obtained from Sigma. Thermo Fisher Scientific was approached for obtaining lysotracker green DND-26 (LTG, #L7526), and Collagen (#A1048301). Phalloidin-iFluor 488 reagent (#Ab176753) was purchased from Abcam. Enhanced Chemiluminescence (ECL, #1705061) was obtained from Biorad. CytoFLEX Sheath Fluid (#B51503) was acquired from Beckman Coulter. Primary antibody for immunofluorescence, p62/SQSTM1 (#NBP1-48320SS) was acquired from Novas Biologicals and Anti-Rabbit IgG Alexa fluor 555 (#A32732) and Anti-Mouse IgG Alexa fluor 488 (#A32723) were procured from Invitrogen. Primary antibodies SQSTM1/p62 (D1Q5S) #39749S, anti-LC3 A/B (D3U4C) #12741S, anti-β-Actin (8H10D10) #3700S, anti-β-tubulin (D3U1W) #86298T, anti-LAMP-1 (D2D1, #9091S) and secondary antibodies viz. anti-mouse (#7076S) and anti-rabbit (#7074P2) were purchased from Cell Signaling Technology. Anti-P-Selectin (CTB201, #sc-8419) was acquired from SantaCruz Biotechnology.

### Animal experiment details

Wistar rats, 6-8 weeks aged with body weight in the range of 120-160g were obtained from Central Animal Facility at BITS Pilani according to the approved protocol no. IAEC/RES/27/01. Animals were housed in polypropylene cages with temperature maintained at 24°C and they were fed ad libitum. Experiments performed in this article were approved by IAEC, BITS Pilani and euthanasia was performed in case of thrombus and organ extraction.

### Isolation of platelet from whole blood

Rats were anaesthetized and blood was collected in acid-citrate-dextrose (ACD) buffer containing eppendorf by retro orbital puncture. Collected blood was distributed in fresh 1.5ml eppendorf and equal volume of modified Tyrode-HEPES buffer was added to it. The contents were centrifuged for 15minutes at 200g for platelet rich plasma (PRP) separation without brake, as that might lead to activation of platelets. The separated PRP was incubated at room temperature for 5minutes. PRP obtained was transferred to 1.5ml eppendorf along with the treatment and incubated for 10minutes. These tubes were re-centrifuged at 800g for 10minutes at 22°C. Obtained pellets were carefully resuspended in modified Tyrode-HEPES buffer.

### *Ex-vivo* exposure of platelets to hypoxia

The culture dishes were precoated with 1% gelatin for adherence of platelets. Platelets re-suspended in modified Tyrode-HEPES buffer were added to these dishes and kept in hypoxia incubator procured from ThermoFisher Company (Model 4131) through project sponsored by LSRB, Deference Research and Development Organization for subjecting to variable oxygen concentrations for 30minutes. The normoxic conditions were maintained at 21% and hypoxic conditions maintained at 10%.^15,16^ Platelets were visualized under inverted microscope before further experimentation.

### Microscopic imaging of isolated and cultured platelets

Upon adhesion, platelets were visualized and imaged under microscope post 30minutes exposure to 21% (normoxic) and 10% (hypoxic) oxygen concentrations using Zeiss Axiocam 105 color microscope. Following this 4% paraformaldehyde treatment was given to the platelets for 10-15minutes for fixation and phalloidin (1:1000) and DAPI (1:1000) stains were added thereafter for platelet characterization.

### Platelet aggregation

Isolated and treated platelets were seeded in 96-well plate and kept under varying oxygen conditions inside the incubator for 30minutes. Post this, readings were taken at 405nm for 10minutes with 2minutes shaking at high speed in the middle using Multiskan sky spectrophotometer procured from Thermoscientific and light absorbance was measured.^7,17^

### Scanning electron microscopy

Culture dishes were coated with 1% gelatin for 1hour and isolated platelets were subjected to variable oxygen concentrations (21% and 10% O_2_). Post 30minutes, platelets were treated with 4% paraformaldehyde for fixation followed by PBS wash. The samples were dehydrated chronologically using 50%, 75% and 100% ethanol for 1minute each and were then air dried. Samples were coated with thin film of chromium in a vacuum coater before visualization using Apreo S of FEI (Thermo fisher) field emission scanning electron microscopy (FESEM).

### Cytotoxicity analysis

Platelets were cultured in gelatin coated 96-well plates along with specific treatments for 30minutes. MTT assay was used to determine cytotoxicity wherein live upon its addition, live cells form formazan crystals which are further dissolved in DMSO, following which absorbance was measured at 570nm and 630nm using Multiskan sky spectrophotometer (Thermo Fisher). Normoxia and hypoxia exposed platelets without any other treatment served as control for calculating relative percentage cellular viability.

### Static adhesion and platelet spreading assay

18mm coverslips were coated with collagen (20ug/ml) and kept at 37℃ as per manufacturer’s protocol. Isolated platelets were distributed on these collagen-coated cover slips and post 30minutes of hypoxia exposure paraformaldehyde was added to fix the cells. After Triton-X treatment, platelets were stained with phalloidin (1:1000) for 30minutes and mounted using glycerol. These slides were further viewed under microscope (Zeiss Apotome).^18^

### Clot retraction assay

PRP was treated with chloroquine for 10minutes and 1U thrombin (platelet activation inducer) was added followed by immediately stirring and sealing the test tube.^19^ They were then exposed to hypoxic and normoxic conditions for 30minutes and retraction of clot was analyzed.

### Lysotracker staining

Washed platelets obtained from whole blood were exposed to 21% (normoxic) and 10% (hypoxic) conditions. 1μM lysotracker dilution was added to the eppendorfs and an incubation of 15-20minutes was given to the samples at 37℃, followed by centrifugation at 400g for 5minutes and resuspension of pellets in buffer. Sample acquisition was done using flow cytometer (Cytoflex, Beckmann Coulter) followed by data analysis by CytExpert software.

### Immunofluorescence

Platelets exposed to variable oxygen conditions for 30minutes. 4% paraformaldehyde was used for fixation and 0.1% Triton-X was added for permeabilization. 2.5% BSA solution was used for blocking so as to avoid any kind of non-specific interaction. After overnight primary antibody incubation, secondary antibody was added and incubated for 2hours post which phalloidin staining was done. The slides were carefully mounted and imaged using 63x/1.4 oil Zeiss LSM880 confocal microscopy. Details of antibodies used for this experiment is referred in ‘chemicals and reagents’ section

### Flow cytometry

Washed platelets were incubated under 10% O_2_ conditions for 30minutes and primary antibody P-Selectin (1:500) was directly added inside the incubator to avoid re-oxygenation. After 30minutes incubation with primary antibody, platelets were centrifuged at 400xg for 5minutes and then Alexa fluor 488 (1:1000) was added to the obtained pellets. It was again incubated for 30minutes and further centrifuged. The obtained pellets were carefully re-suspended in flow buffer, following which samples were acquired by flow cytometer and data was analyzed using CytExpert 2.4.

### Immunoblotting

A cocktail containing platelet lysis buffer (2% NP40, 150mM NaCl, 30mM HEPES, 2mM EDTA) and PMSF (Sigma Aldrich) was added to the samples post 30minutes hypoxia exposure inside the incubator to avoid re-oxygenation. They were kept in 4℃ for 30minutes and then stored in −20℃ before further processing. Samples were snap thawed and pipetted vigorously for cell membrane lysis. After centrifugation at 12000rpm for 20minutes at 4℃, the supernatant was collected. Bradford assay was done to estimate the amount of obtained protein. For sample preparation, 1X buffer was added to loading amount of protein and heat denatured in dry bath at 100℃ for 10minutes. Final samples were loaded in the wells created on polyacrylamide gel and protein separation was carried out with SDS-PAGE technique. The contents of gel were relocated to PVDF membrane using semi dry transfer apparatus (Biorad) followed by blocking of non-specific sites using 5% skimmed milk in TBS. The membranes were probed and re-probed with primary antibody (1:750) overnight at 4℃ and were subjected to secondary antibody incubation for 2hours at room temperature. Horseradish peroxide-conjugated goat antirabbit IgG were used as secondary antibodies. Post washing of any excess antibody from the membranes, enhanced chemiluminescence detection system (Biorad) was used for spotting protein intensities. Densitometric quantification of the detected proteins was performed by ImageJ software by normalization using control.

### Animal exposure to hypoxic conditions

Animals were treated with chloroquine (5mg/kg) 30-45minutes before exposure to hypoxic and normoxic conditions. They were kept in a sealed chamber under experimental conditions for 24 hours with a gas exchange of 5-10minutes after every 3hours to avoid CO_2_ saturation. Blood collected from these animals was used to study parameters related to platelet aggregation.

### Tail bleeding and blood volume analysis

Animals were anaesthetized and a cut was made at 3-5mm distance from the tip of its tail. Blood was collected in an eppendorf containing PBS and time taken for cessation of bleeding was noted.^20^ Volume of collected blood was subsequently measured.

### Flow restriction animal model

Animals were given ketamine (40mg/kg) and xylazine (10mg/kg) according to their body weights. A blood flow restriction model was developed to initiate thrombus formation by proximal ligation of inferior vena cava (IVC) and its lateral tributaries. The formed thrombus was collected after 24 hours of surgery and measured to compare the effects of treatment with that of control.^21^ Post surgery the animals were given gentamycin (25mg/kg) and celecoxib (10mg/kg) treatment in order to avoid infection and pain.

### Histology

Organ samples and thrombus were stored in formalin after extraction for fixation and were outsourced to Advanced Histology Labs, New Delhi for slide preparation. Briefly, the process followed for slide preparation consists of block preparation and subsequent sectioning of the samples which were then placed on slides and stained using haematoxylin and eosin stain. The obtained slides were imaged under microscope (Zeiss).

### Mass spectrometry sample preparation and analysis

Whole blood samples were collected from animal post CQ treatment and 24hours of hypoxia exposure. Platelets were isolated and protein was extracted from the samples using lysis buffer.

Obtained whole cell lysates were outsourced to Valerian Chem Private Limited, New Delhi for mass spectrometry. Briefly, protein per sample was used for digestion and reduced with 5 mM TCEP. Further 50mM iodoacetamide treatment was given for alkylation followed by digestion with Trypsin (1:50, Trypsin/lysate ratio) for 16 h at 37 °C. C18 silica cartridge was used to clean the digests by removing salt accompanied by speed vac for drying. The dried pellet was re-suspended in buffer A (2% acetonitrile, 0.1% formic acid). Experiments were performed on an Easy-nlc-1000 system coupled with an Orbitrap Exploris mass spectrometer. 1ug of peptide sample were loaded on C18 column 15 cm, 3.0μm Acclaim PepMap (Thermo Fisher Scientific) and separated with a 0–40% gradient of buffer B (80% acetonitrile, 0.1%formic acid) at a flow rate of 500 nl/min) and injected for MS analysis. LC gradients were run for 110minutes. MS1 spectra were acquired in the Orbitrap (Max IT = 60ms, AGQ target = 300%; RF Lens = 70%; R=60K, mass range = 375−1500; Profile data). Dynamic exclusion was employed for 30s excluding all charge states for a given precursor. MS2 spectra were collected for top 20 peptides. MS2 (Max IT=60ms, R= 15K, AGC target 100%). All samples were processed and RAW files generated were analyzed with Proteome Discoverer (v2.5) against the Uniprot Rattus database. For dual Sequest and Amanda search, the precursor and fragment mass tolerances were set at 10 ppm and 0.02 Da, respectively. The protease used to generate peptides, i.e. enzyme specificity was set for trypsin/P (cleavage at the C terminus of “K/R: unless followed by “P”). Carbamidomethyl on cysteine as fixed modification and oxidation of methionine and N-terminal acetylation were considered as variable modifications for database search. Both peptide spectrum match and protein false discovery rate were set to 0.01 FDR.

### Statistical analysis

Obtained data was analyzed using GraphPad Prism software version 8.0.1 and plotted applying mean with SD. Statistical determination of role of hypoxia with and without treatment in contrast to control was determined using unpaired t-test. Multiple comparisons were examined using Turkey method with significance defined as p-value ≤ 0.05

## Results

### Hypoxia (10%) induces pro-thrombotic features in platelets

Acute hypoxic conditions are reported to induce a pro-thrombotic state in platelets; however, effect of a physiologically relevant hypoxic state associated with high altitude is poorly understood. To evaluate the same, platelets were isolated by retro-orbital bleeding from female Wistar rats and were immediately cultured *ex vivo* under normoxic (21% O_2_) or hypoxic condition (10% O_2_) following methods mentioned in the ‘materials and methods’. Unlike majority of the existing studies, herein a physiologically relevant oxygen percentage (10% O_2_) that is often experienced at high altitudes in recreational climbers or soldiers was considered over an acute hypoxia.^22,15^ As evident from the phase contrast images and the correlated quantitative bar diagram, an increased aggregation or clumping of platelets was observed under hypoxia **(Fig. 1a)**. In addition, we performed a light transmission aggregometry experiment, as described by Chan et al 2018, which states that light absorbed by platelets is inversely related to the formation of aggregates.^17^ Interestingly, a significantly (*p valu*e 0.0008) reduced light absorbance, putatively attributed to the aggregation of platelets under hypoxic condition was observed in our study **(Fig. 1b)**. Furthermore, abundant cytoskeletal extensions, indicative of activated platelets could be clearly visualized in scanning electron microscopic (SEM) images **(Fig. 1c)**.^23,6^ Moreover, formation of cytoskeletal protrusions after phalloidin staining of platelet actin filaments, and evident from extensive green projections under hypoxia also confirmed an altered platelet homeostasis **(Fig. 1d)**. The above results are indicative of the fact that a non-acute hypoxic condition, like 10% O_2_ can also stimulate a state reflective of activated platelets *ex vivo*. Herein, existing literature suggests that platelets demonstrate an increased adhesive property upon appropriate stimuli to facilitate their clot formation.^23^ Interestingly, a 10% O_2_ exposure also stimulated an increased adhesion of cultured platelets onto collagen coated surfaces **(supplementary figure 1a).** Herein, clot retraction assay is often used as a reproducible, alternate approach to effectively assess platelet activation.^24,18^ Therefore, qualitative estimation of clot formation under hypoxic condition was also performed, which showed that the mesh gets substantially retracted under the hypoxic state, indicative of clotting **(supplementary figure 1b).** Platelet activation is conventionally monitored at the molecular level through analysis of P-Selectin which is generally stored in alpha granules of platelets and are expressed on its membrane surface upon activation. Importantly, P-Selectin protein levels were found to be increased under hypoxic condition, when compared to normoxia, as analyzed through flow cytometry **(Fig. 1e)**. The above results demonstrate the platelet adhesion and activation potential of 10% O_2_ when cultured *ex vivo*; however, the key molecular signals driving it remains to be further elucidated.

**Figure 1:**
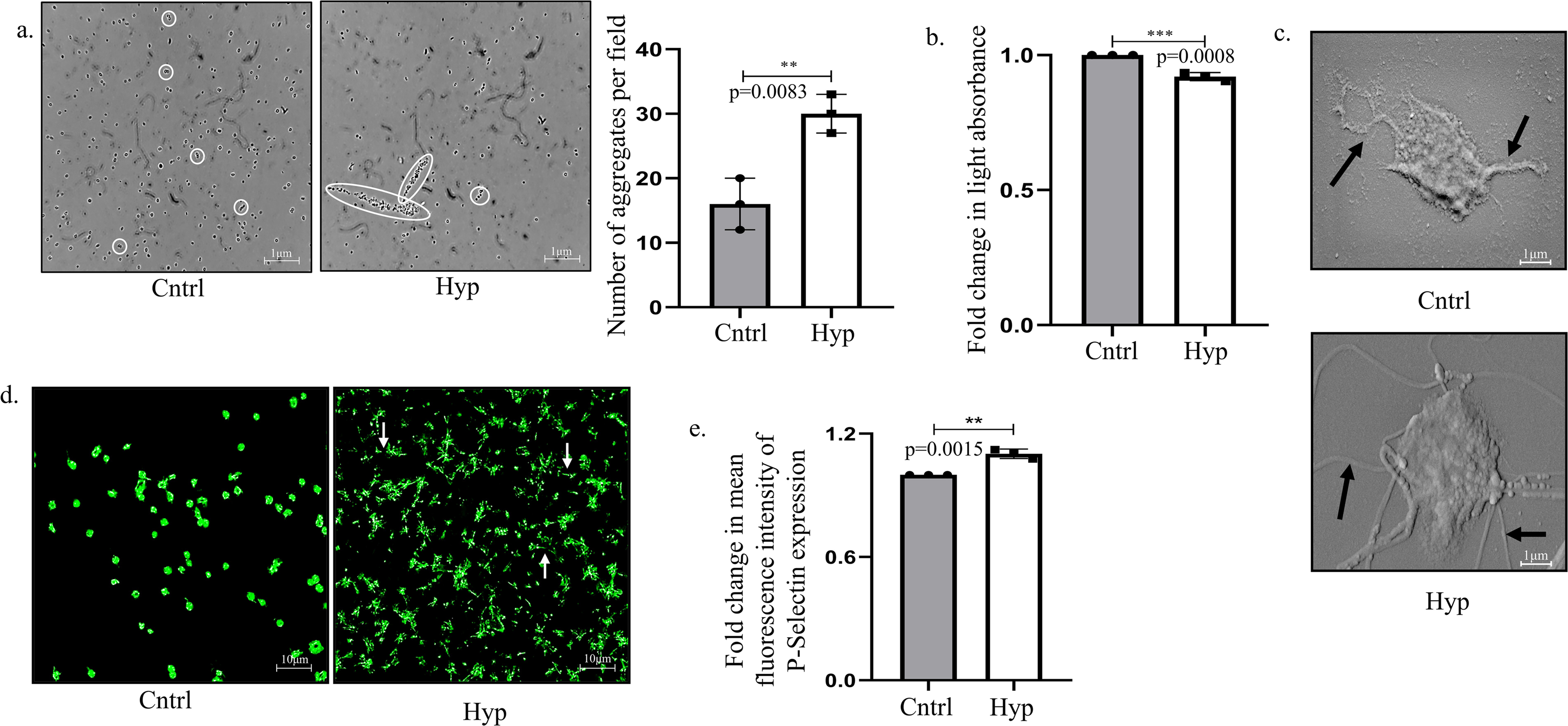
Hypoxia (10%) induces pro-thrombotic feature in platelets. **(a)** Phase-contrast microscopic images and graphical representation of platelet aggregates under normoxic (Cntrl; 21% oxygen) or hypoxic condition (Hyp; 10% oxygen; 30minutes exposure) (Scale bar: 1μm). **(b)** Graphical representation of light transmission aggregometry showing fold change in light absorbance owing to exposure to normoxic or hypoxic state. **(c)** Scanning electron microscopy images of platelets showing cytoskeletal extensions from platelet surface (denoted by black arrows) under hypoxia compared to normoxia. **(d)** Phalloidin staining of cells depicting filamentous bridges (denoted by white arrows) formed between platelets under low oxygen conditions (10% oxygen for 30minutes) (Scale bar: 10μm). **(e)** Mean fluorescence intensity of P-Selectin as analyzed by flow cytometry after 30minutes exposure to 21% oxygen (Cntrl) or 10% oxygen (Hyp). **p<0.01, ***p<0.001.

### Exposure of platelets to 10% hypoxia triggers autophagy

Recent literature indicates that cellular structures associated with autophagy and its degradative function are present in platelets and may play a key role in its associated functions like clotting, inflammation or immunity. Importantly, though autophagy is well-studied in nucleated cells, its function in anucleate platelet cells is way less recognized. Moreover, since platelets are cells without a nuclear machinery to continuously synthesize new mRNAs, tracking autophagy in these atypical cells is tricky. We hence followed methods from Madhu M Ouseph et.al. 2015 who proved that a reduction in LC3-II can serve as a clear indicator of autophagy in both activated (with thrombin) murine and human platelets.^8^ On a similar note, in a separate study Tzu Yin Lee et al 2021 reported that the adapter protein SQSTM1/p62 undergoes degradation in activated human platelets and a reduced p62 protein conventionally serves as a marker of autophagy.^25^ In corroboration to above, we also observed a decline in both LC3-II and p62 under 10% O_2_ confirming an activation of autophagy **(Fig. 2a).** In this context, Ouseph et al also proposed that an active proteolysis of LC3-II in acidic compartments results in its reduction. There was a simultaneous decrease in fluorescence intensity of p62 protein in platelets exposed to hypoxic environment as compared to normoxia **(Fig. 2b)**. Interestingly, our flow cytometric analysis revealed a significant increase in lysotracker fluorescence post hypoxia exposure further validating a putatively active autophagic flux in the platelets **(Fig. 2c)**. Herein, the lysosomes serve as degradation hub for autophagosomal or endocytic vesicles and the lysosomal associated membrane protein 1/2 (LAMP1/2) is known to be distributed amongst these heterogenous endocytic, autophagic and endolysosomal structures;^26^ thus a reduction in protein level of LAMP1/2 can often serve as an indicator of active autophagic flux.^27^ As published by Monaci et. al. in 2022, hypoxia increases acidity in dendritic cells causing increase in lysotracker fluorescence along with reduced LAMP1 intensity.^28^ Importantly, we also observed a reduced fluorescence intensity of LAMP1 in platelets exposed to 10% oxygen concentration along with decrease in LAMP2A protein expression **(Fig. 2d and 2e)**.^29^

**Figure 2:**
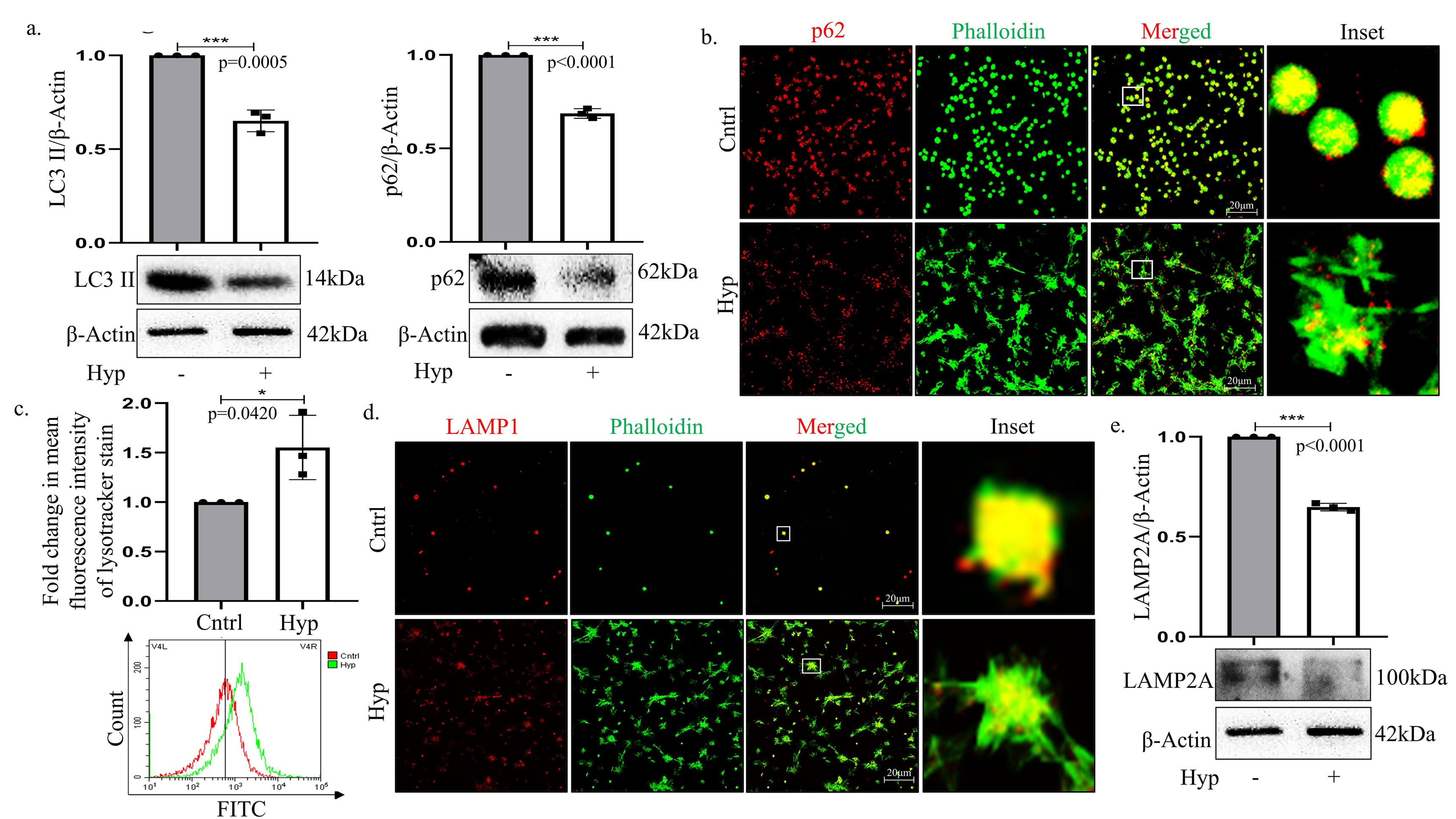
Hypoxia induces autophagy in platelets. **(a)** Immunoblot of autophagy associated proteins like LC3II and p62 in platelets exposed to normoxia or hypoxia for 30minutes. **(b)** Immunofluorescence of p62 protein (Red) in platelets upon subjecting them to hypoxia (30minutes) compared to normoxia. Cells were co-stained with phalloidin (Green) (Scale bar: 20μm). **(c)** Graphical representation of lysosomes in platelets under hypoxia as measured with lysotracker by flow cytometry. **(d)** Immunofluorescence of LAMP1 (Red) in platelets exposed to varying oxygen conditions-Normoxia (Cntrl; 21%) or Hypoxia (Hyp; 10%) for 30minutes. Cells were co-stained with phalloidin (Green) (Scale bar: 20μm). Inset depicts zoomed image. **(e)** Immunoblot of LAMP2A expressed by platelets when exposed to hypoxia for 30minutes. *p<0.05, ***p<0.001.

### Chloroquine inhibits hypoxia induced autophagy in platelets *ex vivo*

Given that autophagy has earlier been described for its role in platelet activation and our current findings also indicate that hypoxia stimulates autophagy in platelets, we therefore decided to inhibit the homeostatic process with existing drugs. In this regard, the current practice is to impair autophagy by blocking the lysosomal degradation process with compounds like, bafilomycin (Baf) or chloroquine (CQ) or protease inhibitors.^30^ Amongst the above, CQ and HCQ (hydroxychloroquine) are FDA approved, have been widely repurposed and are hence the primary choice for use in clinics in various diseases. CQ disrupts the binding of autophagosomes with lysosomes and can have non-autophagic effects as well like-disorganization of the cytoskeleton or Golgi/endo-lysosomal organization.^30^ Therefore, we exposed platelets *ex vivo* to CQ under both normoxic or hypoxic condition and analyzed its cytotoxic and autophagy inhibitory potential on platelets. As evident from the **supplementary figure 1c** the specific dose of CQ (2μM) used for subsequent study had a minimal cytotoxic effect on the platelets. Importantly and as expected, CQ treatment resulted in an increased accumulation of proteins like LAMP2A, LC3II and p62 in platelets as compared to only hypoxic condition, as analyzed by immunoblot **(Fig. 3a and 3b)**. Furthermore, immunofluorescence analysis also showed an accumulation of red fluorescence representative of LAMP1 and p62 accumulation in platelets exposed to CQ implicating a buildup of vesicles positive for these markers due to disrupted cytoskeletal trafficking **(Fig. 3c and supplementary figure 1d)**. In addition, exposure of platelets to lysotracker that is known to stain acidic lysosomal compartments also depicted reduced fluorescence. CQ has been earlier reported to alter the lysosomal pH thus interfering with autophagosome-lysosome fusion;^31^ the same was found to be true for atypical cells like platelets as well **(Fig. 3d)**. Therefore, we hypothesized that the FDA-approved drug CQ can be an excellent choice to inhibit autophagy in platelets to modify their function under hypoxic condition, which was further explored in this study.

**Figure 3:**
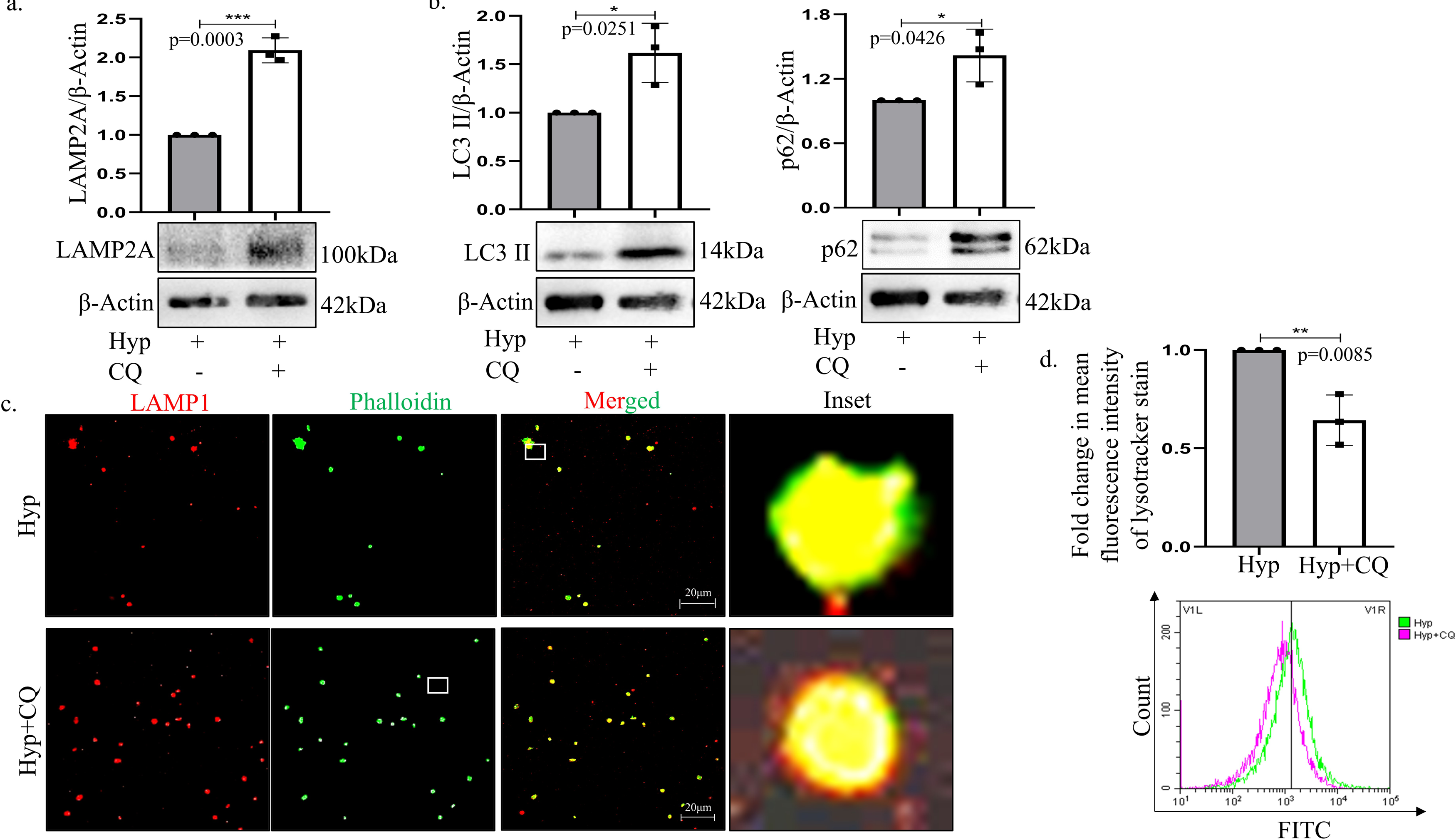
Chloroquine inhibits platelet activation. **(a)** Immunoblot of LAMP2A in platelets exposed to either hypoxia (10% oxygen; Hyp) or hypoxia plus chloroquine (CQ; 2μM) for 30minutes. **(b)** Immunoblot of autophagy associated markers-LC3II and p62 in platelets treated with or without CQ under hypoxia for 30minutes. **(c)** Immunofluorescence of LAMP1 (Red) in platelets subjected to CQ and hypoxia post 30minutes exposure. Cells were co-stained with phalloidin (Green) (Scale bar: 20μm). **(d)** Fold change in lysotracker staining in platelets when exposed to hypoxia in presence or absence of CQ. *p<0.05, **p<0.01, ***p<0.001.

### Chloroquine attenuates hypoxia-induced platelet activation *ex vivo*

To understand the effect of CQ on platelet function, we pre-treated the platelets with the drug followed by exposure to hypoxia. Interestingly, analysis of platelet aggregation under phase-contrast microscope revealed a significant decrease in the cellular aggregate formation in presence of CQ **(Fig. 4a)**. On a similar note, light transmission aggregometry also demonstrated an increase in light absorbance by platelets upon treatment with CQ reflective of a decreased platelet aggregation **(Fig. 4b)**. In addition, phalloidin staining showed a decreased cytoskeleton-mediated platelet-platelet interaction upon exposure to CQ suggesting that autophagy inhibition can have a negative impact on platelet aggregative properties induced by hypoxia **(Fig. 4c)**. Platelets upon activation reorganizes its actin cytoskeleton resulting in formation of spiky filamentous structures.^32^ Interestingly, scanning electron microscopy clearly showed a drastic reduction of hypoxia induced cytoskeletal extensions upon autophagy inhibition with CQ **(Fig. 4d)**. Further, the clot retraction assay post CQ treatment depicted an expansion of mesh supporting the potential of CQ in reducing unnecessary clot formation **(supplementary figure 1e).** Proteins stored in platelet alpha-granules play a significant role in identification of platelet activation status, and P-Selectin is the major protein secreted from these storage granules and represents an activated state of platelets. As expected, there was an attenuation of P-Selectin levels upon autophagy inhibition **(Fig. 4e)**. Thus, from the above experiments, it can be inferred that autophagy plays a positive role in platelet activation under hypoxic condition, and its inhibition can potentially lower clot formation.

**Figure 4:**
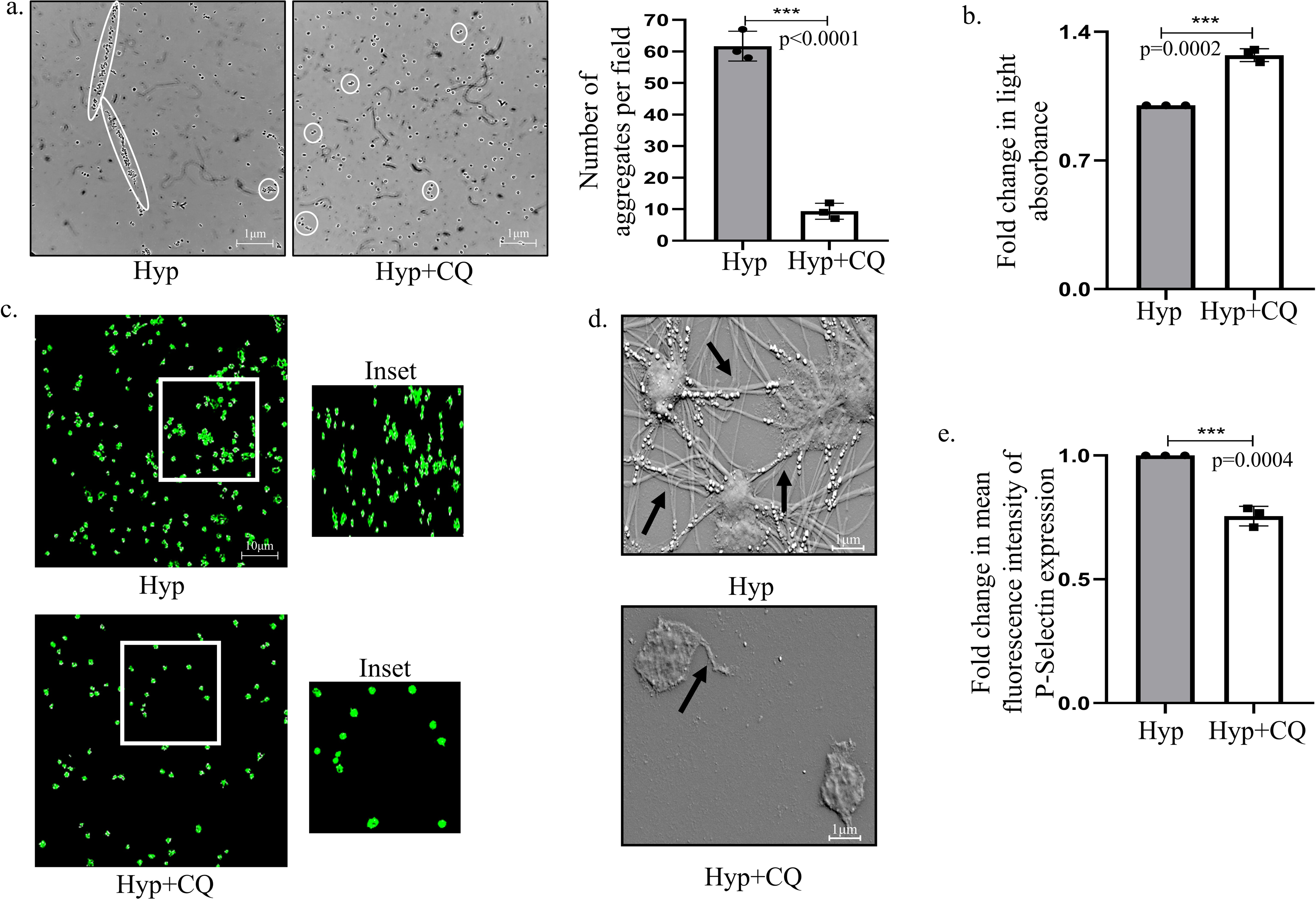
Chloroquine attenuates hypoxia-induced platelet activation *ex vivo*. **(a)** Images and bar graph showing platelet aggregates under hypoxia (10% oxygen; 30minutes) in presence or absence of CQ (2μM) (Scale bar: 1μm). **(b)** Graphical representation of light absorbed by platelets under hypoxia in presence or absence of CQ. **(c)** Phalloidin staining of cytoskeletal filaments in platelets exposed to CQ under hypoxia (Scale bar: 10μm). **(d)** Scanning electron microscopy of platelets after CQ treatment under hypoxia. Black arrows represent the filamentous processes observed. **(e)** Graphical representation of fluorescence intensity of platelet activation mark-P-Selectin under hypoxia in presence or absence of CQ, as analyzed through flow cytometry. ***p<0.001.

### Chloroquine inhibits hypoxia-induced platelet activation *in vivo*

To simulate a closer life mimicking scenario we further inhibited autophagy *in-vivo*. Physiologically, the conventional platelet functions are triggered upon stress conditions induced either by the environment or vascular injury. Herein, hypoxia is reported to act as one such inducer increasing platelet count and activation under *in-vivo* conditions.^31^ Thus, to explore the potential of CQ under physiologically relevant hypoxic condition, animals were pre-treated with CQ and subjected to 10% O_2_ for 24 hours in a hypoxia chamber, constructed in-house for hypoxia treatment.^21,33^ Interestingly, an increased bleeding time (tail bleeding experiment) and enhanced blood volume was observed with CQ treatment as compared to hypoxic control animals **(Fig. 5a and 5b)** conclusive of the fact that time required for clot formation is increasing with CQ treatment under 10% O_2_. Further, blood samples were collected from these animals to explore platelet activation. To achieve this, light transmission aggregometry was carried out and results showed an increase in light absorbance indicative of decreased platelet aggregates with CQ under hypoxia **(Fig. 5c)**. Furthermore, P-Selectin also showed a reduced expression with CQ under 10% O_2_ confirming a decrease in platelet activation *in-vivo* **(Fig. 5d)**. In accordance to the above results, an increase in p62 and LC3 expression was found in platelet samples isolated post hypoxia and CQ treatment confirming an autophagy inhibition **(Fig. 5e)**. In addition, whole cell proteomic analysis (from platelet isolated from rats) was also performed to have a profile of proteins de-regulated under only hypoxia, or CQ with hypoxia exposure. Importantly, in corroboration to our earlier findings, a de-regulation in expression pattern of proteins associated with platelet adhesion, activation and aggregation, and also autophagy was observed under hypoxia; a list of some of the proteins de-regulated and involved in the above processes is presented in the **supplementary figure 2a.** Interestingly, for example, integrin linked protein kinase (ILK) which is known to play a major role in platelet function, initiation of thrombus formation, and is also positively implicated in autophagy, was found to be expressed in hypoxia but was absent in CQ treated group.^34,35^ Further research is required in this direction to dissect the role of such individual proteins in the above context. Finally, to ascertain that CQ dose administered has minimal cytotoxic effect on cellular tissues, histological sectioning of liver and kidney was performed after CQ treatment. Importantly, it did not show any significant aberrations or cytotoxic effect suggesting that CQ, at the stipulated dose was well tolerated and had minimal effect on the tissue integrity **(supplementary figure 2b and 2c).** All the above observations from both ex*-vivo* and *in vivo* system suggest that CQ can be used as a putative alternative to compromise platelets’ activity and thus thrombotic clot formation through inhibition of autophagy under low oxygen conditions.

**Figure 5:**
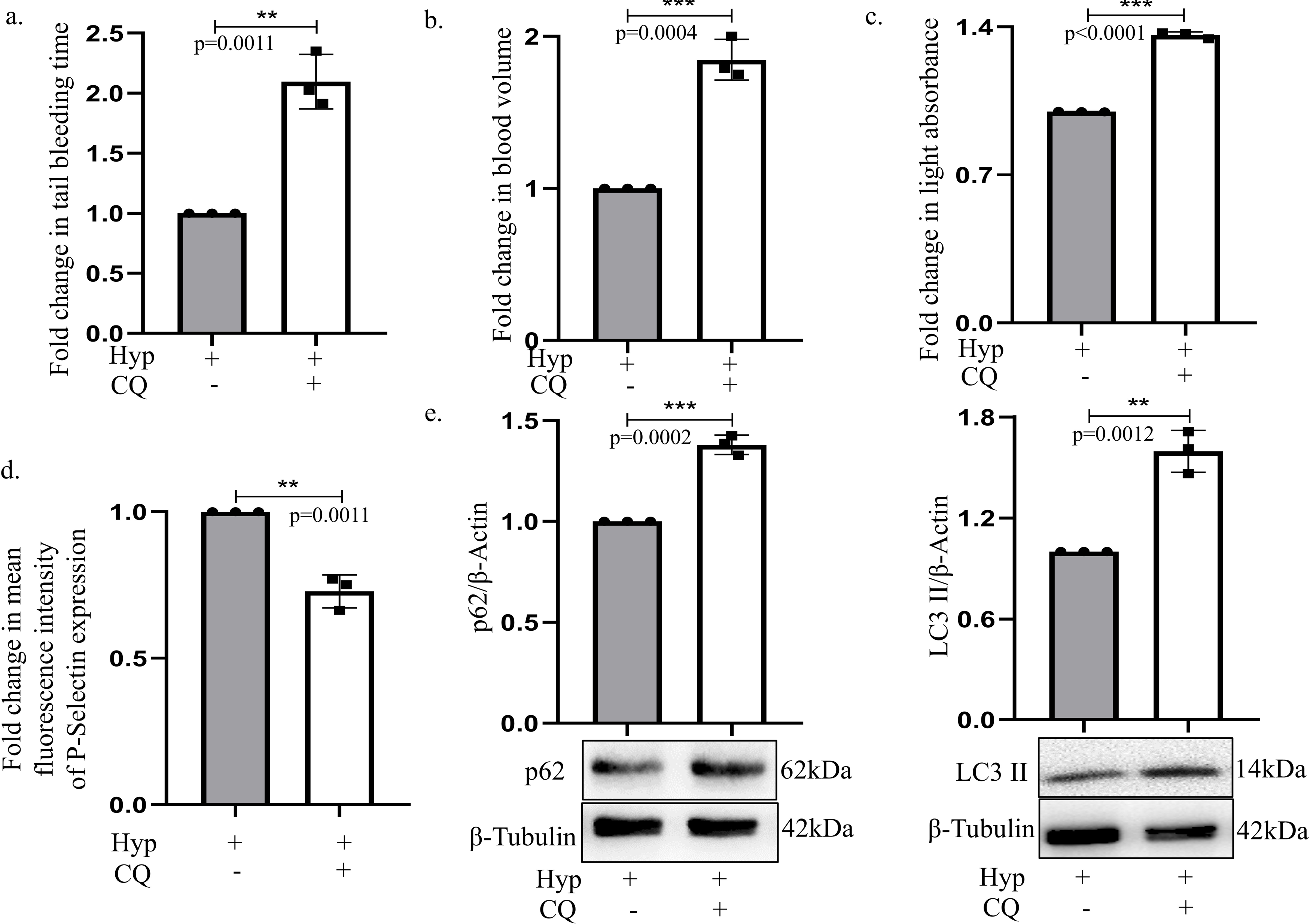
Chloroquine inhibits platelet activation *in vivo*. **(a)** Graphical representation of fold change in bleeding time or time taken for clot to form in CQ pre-treated (5mg/kg body weight; intraperitoneal; 30minutes) animals exposed to hypoxia (10% oxygen; 24hours). **(b)** Graphical representation showing fold change in blood volume collected from CQ-treated animals under hypoxic condition. **(c)** Graphical representation of light transmission aggregometry based fold change in light absorbance in platelets obtained from *in-vivo* conditions wherein animals were subjected to hypoxia with or without CQ treatment. **(d)** Graph representing fold change in fluorescence intensity of P-Selectin in rats exposed to hypoxia (24hours) pre-injected with intra-peritoneal CQ. **(e)** Immunoblot of p62 and LC3II expressed by *in-vivo* derived platelets when rats were subjected to hypoxia for 24 hours post intra-peritoneal CQ injection. **p<0.01, ***p<0.001.

### Chloroquine causes decrease in thrombus formation in flow restriction model

To finally confirm our results obtained above, a flow restriction model was developed by ligating the inferior vena cava (IVC) and its lateral tributaries so as to initiate thrombus formation in rats. This ligation has already been proven in earlier studies as a source of rapid venous thrombi generator which can induce thrombus formation as early as 15minutes post-surgery.^36^ Herein, the ligation of blood vessels leads to restricted blood flow causing platelets to get activated and form thrombus due to their aggregation. This surgery that results in deep vein thrombus was performed to simulate a thrombosis that might occur *in vivo* at high altitude.^21^ Interestingly, if CQ was administered 30minutes prior to surgery, a reduced thrombus size was observed in surgical animals, compared to non-treated animals, as evident from the thrombus image (**Fig. 6a).** A clean IVC was observed in SHAM control animals when compared to ligated animals confirming that thrombus formation is initiated by surgical restriction of blood flow and not by abdominal incision **(Fig. 6b).** Upon measurement of thrombus size and wet weight, a reduction in both the parameters was evident upon surgical restriction coupled to CQ exposure compared to control or non-inhibition **(Fig. 6c and 6d).** Finally, histology of the isolated thrombus confirmed a blockage in blood vessel, as indicated by black arrow in untreated animals, whereas a clearer blood vessel was observed in CQ treated animals **(Fig. 6e)**. Thus, these results affirm that CQ effectively inhibits autophagy in flow-restricted animal models as well, causing decrease in thrombus size and hence can be a potential arsenal attenuating thrombosis.

**Figure 6:**
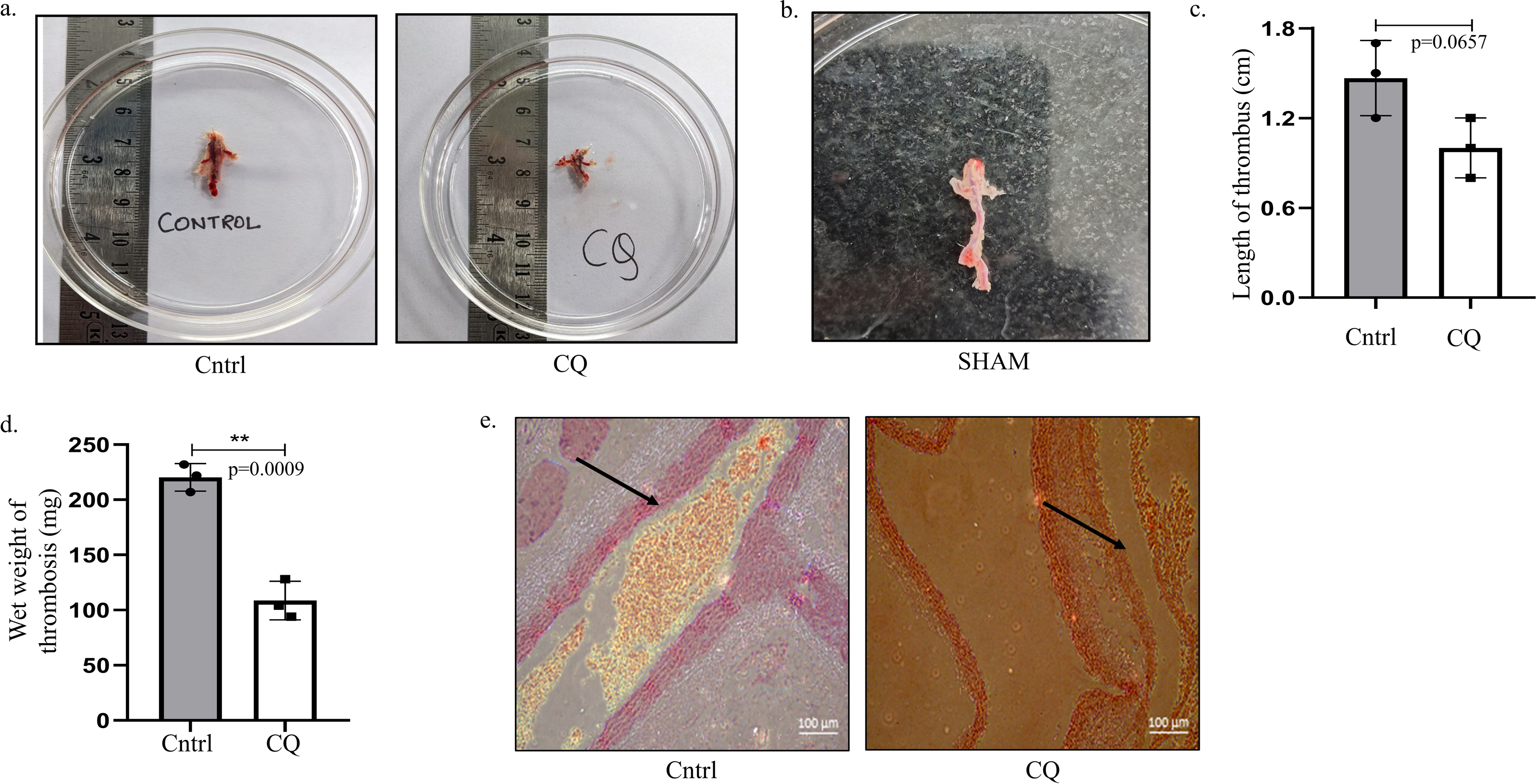
Autophagy inhibition causes decrease in thrombus formation in flow restriction model. **(a)** Thrombus images collected 24hours post ligation of inferior vena cava (IVC) with or without CQ pre-treatment (5mg/kg body weight; intraperitoneal; 30minutes). **(b)** Inferior vena cava region collected from SHAM animal (non-ligated) model post 24 hours of surgery showing absence of thrombus in it. **(c)** Graphical representation of thrombus length (cm) obtained post-surgery from CQ treated or untreated animals. **(d)** Graphical representation of wet weight of isolated thrombus from CQ-treated or untreated animals. **(e)** Histology of IVC showing blood vessel blockage in ligated but CQ untreated animals, compared to clean vessel observed with CQ treatment post 24hour surgery (Scale: 100μm). **p<0.01.

## Discussion

Autophagy is a cellular homeostatic mechanism that regulates energy distribution in our body by engulfing and eliminating damaged and unwanted organelles and/or proteins and lipids of the cells. Its importance during starvation, disease progression and in other stress response is quite well known. Interestingly, a recent article by Feng et.al. reported presence of autophagy machinery and its key role in platelet function.^37^ Therefore, platelets-despite being cellular fragments shredded during megakaryocyte maturation, possessing limited life span, restricted cellular machinery to sustain, have mechanisms like autophagy that can regulate homeostasis. In accordance to above, in our study, we observed that autophagy is up-regulated and plays a significant role in governing platelet function upon exposure to physiologically relevant hypoxic condition. A low oxygen condition triggered translation of autophagic mRNAs presumably pre-present in resting platelets, leading to platelet activation and turning on their aggregation-resulting in a thrombotic condition. We evaluated the involvement of autophagy in medicating this process and validated it in both *ex-vivo* and *in-vivo* conditions under the low oxygen conditions. Earlier, Lee et.al. reported the link between the two processes, however, our group is arguably the first to show the association between the cellular homeostatic process autophagy and the hemostatic process under 10% oxygen levels.^25^ These results are of immense relevance as various physical activities like recreational climbing, training, pregnancy and sudden movement of low-landers to high altitude are reported to induce such physiological low oxygen conditions pre-disposing an apparently healthy individual to an abrupt state of medical attention. While hypoxia regulating proteins have been explored as therapeutic target against such thrombotic tendency, we for the first time propose the use of chloroquine (CQ) as a therapeutic alternative.^12^ CQ is an FDA approved drug currently marketed as an anti-malarial agent and is presently undergoing clinical trial along with hydroxychloroquine as an autophagy inhibitor for tumor treatment.^30^ Its role as autophagy inhibitor in platelets and subsequently in modifying thrombosis under physiologically relevant hypoxic condition has not been evaluated earlier. Herein, we show through *ex vivo*, *in vivo* and surgical rat model that CQ can effectively inhibit autophagy and platelet function as illustrated in graphical abstract. In addition, total proteomic analysis from CQ exposed rats also confirmed the platelet inhibitory activity of CQ. Importantly, the conventional therapeutic strategy for thrombosis includes the use of marketed formulations such as anti-thrombotic agents-warfarin, dabigatran, edoxaban, aspirin, etc. which often requires long term administration and also carry the disadvantage of blood thinning that might elevate blood loss post injury. Therefore, our study is a therapeutic breakthrough depicting not only the role of autophagy in governing platelet functionality under clinically relevant hypoxic conditions but it also investigates the therapeutic effects of CQ - an already approved drug that can therefore be re-purposed saving time, money and efforts required towards introduction of a novel therapy. In addition, autophagy being an integral part of platelet life span starting from its differentiation from megakaryocytes to its activation ^38,39,40,41^, the fundamentals of how specific autophagic proteins regulate platelet activation shall be very interesting to dissect in the future years to come.

## Acknowledgement

The authors express their gratitude towards Birla Institute of Technology and Sciences (BITS) Pilani, Pilani campus for providing infrastructural facilities and we are extremely thankful to Institutional Animal Ethics Committee (IAEC), BITS Pilani for approving and supporting our animal studies. We would also like to acknowledge Mr. Suman Kumar and Mr. Om Prakash, operators of confocal microscope and scanning electron microscope facilities, respectively for their generous assistance in completing our studies. The graphical abstract was created using BioRender.com. The authors confirm contribution to this paper as follows: Study conception and design: Prof. Rajdeep Chowdhury, Prof. Syamantak Majumder, Prof. Shibasish Chowdhury; experimental execution, data collection, interpretation and analysis: Propanna Bandyopadhyay, Yash T. Katakia; draft manuscript preparation: Prof. Sudeshna Mukherjee, Prof. Rajdeep Chowdhury, Propanna Bandyopadhyay, Yash T. Katakia.

## Sources of Funding

This study was funded by Life Science Research Board (LSRB), Defence Research and Development Organization, project of Prof. Rajdeep Chowdhury (O/o DG (TM)/81/484222/LSRB-351/PEEE&BS/2019). Propanna Bandyopadhyay acknowledges LSRB and BITS Pilani for providing student fellowship.

## Disclosures

### a. Conflict-of-interest disclosure

There is no conflict of interest between the authors of this research article and the funding agency.

### b. Ethics Approval Statement

Animal studies conducted for the experiments mentioned in this manuscript were approved by Central Animal Facility at Birla Institute of Technology and Science (BITS)-Pilani according to the approved protocol no. IAEC/RES/27/01

## Abbreviations

O_2_: Oxygen
CQ: Chloroquine
IVC: Inferior Vena Cava
ROS: Reactive Oxygen Species
DVT: Deep Vein Thrombosis
VTE: Venous Thromboembolism
ACD: Acid-Citrate-Dextrose
SEM: Scanning Electron Microscopy
CO_2_: Carbon Dioxide
p62: Ubiquitin-binding protein (Sequestosome-1, SQSTM-1)
LC3: Light Chain 3 (Microtubule-associated protein 1A/1B)
TCEP: Tris (2-carboxyethyl) phosphine
MS: Mass Spectrometry
LAMP: Lysosomal Associated Membrane Protein Baf – Bafilomycin
HCQ: Hydroxychloroquine
ILK: Integrin linked kinase

---

**Supplementary figure 1: (a)** Static adhesion and platelet spreading assay showing adhesiveness of phalloidin stained platelets on collagen coated cover slips under normoxic (Cntrl; 21% oxygen) or hypoxic (Hyp; 10% oxygen) conditions. **(b)** Pictorial representation of clot retraction assay under treatment conditions of 21% oxygen (Cntrl) or 10% oxygen (Hyp). **(c)** Chloroquine (CQ) dosing in platelets under 21% oxygen (Cntrl) condition as measured by MTT assay. **(d)** Immunofluorescence of p62 protein (Red) in platelets treated with CQ under hypoxic condition (10% oxygen). Cells were co-stained with phalloidin (Green) (Scale bar: 20μm). **(e)** Pictorial representation of clot retraction assay performed post CQ treatment in platelets under hypoxic condition (10% oxygen). ***p<0.001.

**Supplementary figure 2: (a)** Bar graph of platelet proteomic analysis using mass spectrometry of isolated platelets obtained from animals treated with 10% oxygen (Hyp) or 10% oxygen along with CQ (Hyp+CQ) for 24 hours. **(b)** Histology of liver tissue collected from animals subjected to normoxia (Cntrl, 21% oxygen) or hypoxia (Hyp, 10% oxygen) plus CQ treated animals (Hyp+CQ). The images are labelled with black arrow for central vein and orange arrow for hepatocytes. **(c)** Histology of kidney obtained from animals post normoxia (Cntrl, 21% oxygen), hypoxia (Hyp, 10% oxygen) and CQ along with hypoxia exposure (Hyp+CQ). Images are marked with black arrow for glomerulus, blue arrow depicting proximal convoluted tubule (PCT) and green arrow showing distal convoluted tubule (DCT).

## Notes

### Competing Interest Statement

The authors have declared no competing interest.

